# Pseudo-Central Retinal Blood Vessel in Pig, Sheep and Goat

**DOI:** 10.64898/2026.11.18.767657

**Authors:** Xinyue Wang, Susannah Waxman, Ian A. Sigal

## Abstract

Humans and non-human primates have central retinal blood vessels (CRBVs) through their optic nerve head (ONH). The collagen of pig, sheep and goat ONHs also forms structures that look like CRBVs. We aimed to determine whether the CRBV-like structures in pig, sheep and goat are true CRBVs or if they should instead be called pseudo-CRBVs.

Coronal sections through the ONHs of human, monkey, pig, sheep and goat eyes were imaged for collagen. We then used DAPI, multicolor DiOlistics and GFAP immunostaining to visualize nuclei, astrocytes and other ONH constituents. CRBV-like structures were classified into three types: true CRBVs if the structure was only a blood vessel; fully pseudo-CRBV if the evidence indicated only neural tissues or cells other than erythrocytes within; or partial pseudo-CRBV if some of the region appeared as a vessel and other parts did not.

All eyes exhibited CRBV-like structures in the collagen. In human and monkey eyes they were all true CRBVs. Conversely, the CRBV-like structures were fully or partially pseudo-CRBVs in 80% (4/5) of pig, 50% (3/6) of sheep and 100% (2/2) of goat eyes.

In majority of pig, sheep and goat eyes, CRBV-like structures were actually pseudo-CRBVs. Often, what appeared like a vessel lumen, was instead filled by neural tissues reminiscent of pores. The mechanisms that lead to CRBV-like structures that are not actually CRBVs in these three species, but not in humans and monkeys, remain unknown.

## Introduction

Human and monkey eyes have central retinal blood vessels (CRBVs) through their optic nerve heads (ONH) (Albrecht May, 2008; Anderson, 1970, 1969; Hayreh, 1996). When visualized using techniques optimized for collagen, the ONHs of pig, sheep and goat also exhibit what looks like CRBVs: thick collagen structures with an opening suggestive of a vessel lumen (Figure 1) (Brown et al., 2007; Campbell et al., 2015; Feola et al., 2017; Jan and Sigal, 2018; Waxman et al., 2025; Yang et al., 2018). Our goal was to determine if the CRBV-like structures in the ONHs of pig, sheep and goat are actually CRBVs. We propound that if the collagen opening of a CRBV-like structure contains neural tissue such as retinal ganglion cell axons or astrocytes, and/or cells other than erythrocytes, then it should instead be considered a pseudo-CRBV. Whether the CRBV-like structures in pig, sheep and goat are CRBVs or pseudo-CRBVs will have implications for understanding of ONH physiology and should be considered when thinking of these species’ similarities and differences with humans and monkeys.

**Figure 1:**
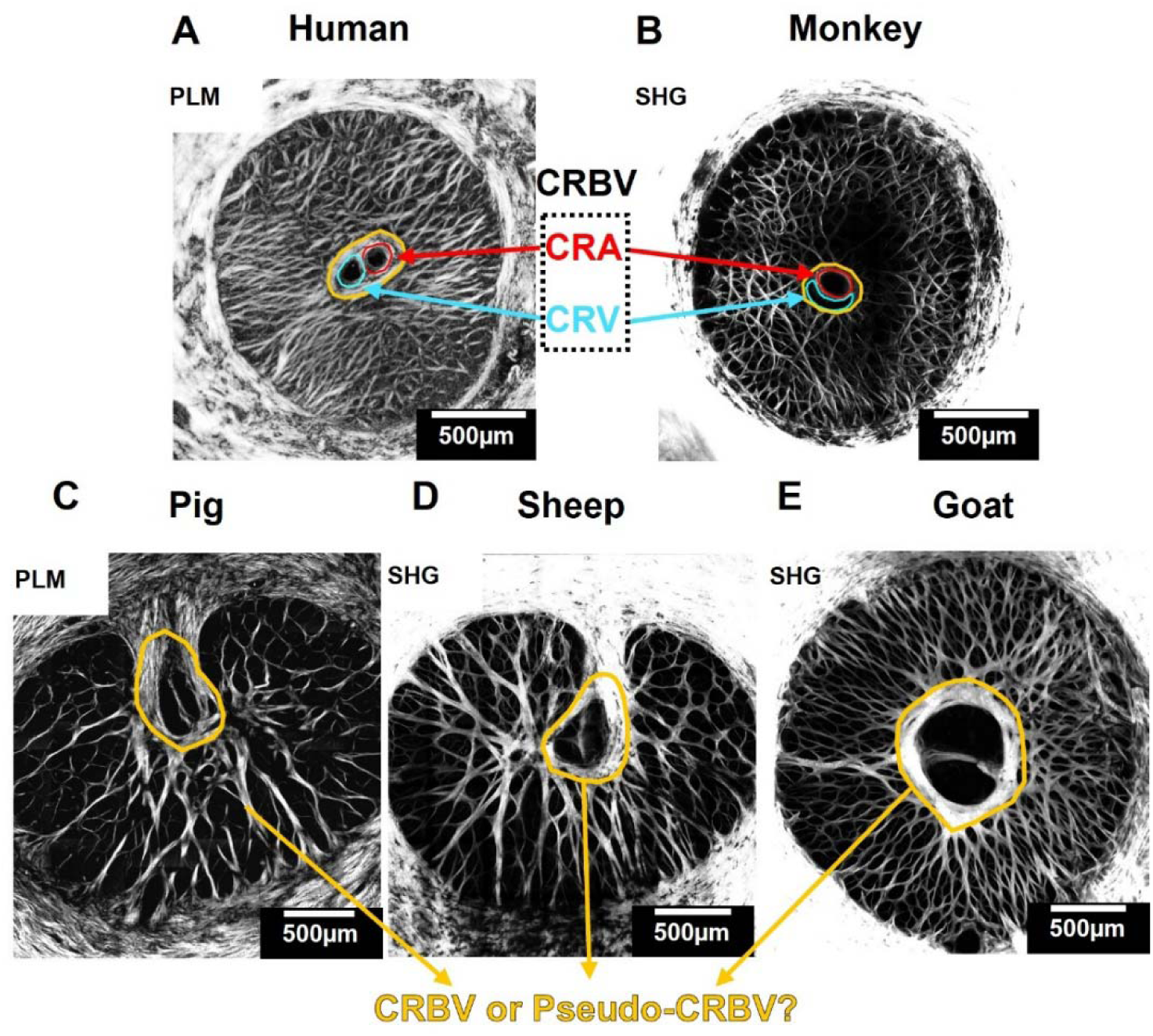
The CRBV-like structures in Pig, Sheep and Goat Look Similar with Human and Monkey. A-B) PLM and SHG images of human and monkey LCs showing collagen network, respectively. Both have CRBVs, well-validated structure in primates, outlined in orange. C-E) SHG and PLM images of the pig, sheep and goat LCs, respectively. A similar central collagenous region is also present in these species, as outlined in orange. This may lead to the conclusion that these structures also contain vessels. PLM: Polarized Light Microscopy, SHG: second harmonic generation, CRA: Central Retinal Artery, CRV: Central Retinal Vein, CRBV: Central Retinal Blood Vessel, LC: Lamina Cribrosa.

## Methods

**Our overall strategy was as follows:** we prepared coronal tissue sections through the ONH at the lamina cribrosa (LC) level of human, monkey, pig, sheep and goat eyes. The sections were imaged using techniques optimized to visualize collagen. Based on these images we confirmed that every eye exhibited a CRBV-like structure. We then used labeling and fluorescence microscopy to visualize nuclei, astrocytes and other non-collagen components. We then classified the CRBV-like structures based on whether these components were present or not into fully pseudo-CRBVs, partial pseudo-CRBVs and CRBVs. Details of each step and definition are provided below.

### 2.1 Eye Procurement and Preparation

This study was conducted in accordance with the tenets of the Declaration of Helsinki, the Health Insurance Portability and Accountability Act, and the Association of Research in Vision and Ophthalmology (ARVO) ’s statement for the use of animals in ophthalmic and vision research. Human, monkey, pig, sheep and goat eyes were utilized for experiments (N= 1, 2, 6, 8 and 2 eyes, respectively). All eyes were ostensibly healthy, without known abnormalities and processed within 24 hours of death as described before (Gogola et al., 2018). The normal human donor eye was obtained through the Center for Organ Recovery and Education of Pennsylvania originally for a different project on cornea. The eye was stripped of all information that could potentially be used to identify the patient. The eye was immersion fixed at 0 mmHg, and later an 11mm trephine used to isolate the ONH. Two Rhesus macaque monkey eyes were obtained from collaborators from animals that had been used for studies not involving the eyes within 30 minutes of death. Pig, sheep, and goat eyes were acquired from a local abattoir within 1 hour of death. No live animals were used in this work. These were processed as the human and monkey eyes: the extraocular tissues were carefully removed and the ONHs were isolated with a trephine, then fixed in 4% PFA for 2–24 hours and held in PBS at 4°C.

The number of human and monkey eyes used in this study was small because these species have already been well-characterized in the literature, and it is not our intent to repeat those studies. Our goal was to produce comparable images of those species that help readers understand why the structures in question in pig, sheep and goat are considered CRBV-like. Although the number of sheep, pig and goat eyes is also small compared with clinical studies, we will show that it is sufficient to satisfy our goal and show that sometimes, CRBV-like structures are Pseudo-CRBVs. Larger sample sizes would be needed to determine the fraction of eyes that exhibit pseudo-CRBVs more precisely in a given population, but we argue that this is secondary to first establishing that pseudo-CRBVs are real.

### 2.2 Sectioning

#### 2.2.1 Cryosectioning

After fixation, 5 pig and 6 sheep ONHs were immersed in a 30% sucrose solution for cryoprotection and embedded in Optimal Cutting Temperature (OCT) compound (Fisher Scientific, Pittsburgh, PA) before freezing in liquid nitrogen (−196 °C). Frozen blocks of ONH tissue were then cryosectioned coronally at 16 µm thickness (Leica CM3050 S, Leica Biosystems Inc., Buffalo Grove, IL).

#### 2.2.2 Vibratome sectioning

The rest of the ONHs were also fixed, and then embedded in low melting temperature 3% agarose (Sigma, A9045), secured to the stage of a vibratome (Leica, VT1200S) with cyanoacrylate glue, and immersed in ice-cold PBS. Coronal slices were cut at 150μm-thickness. Slices were stored in PBS at 4°C for up to 48 hours between sectioning and labeling.

A strength of vibratome slices is that it is possible to visualize 3D structures. This, however, requires 3D techniques that are slower and more complex than for visualizing 2D structures in cryosections.

### 2.3 Labeling and Staining

For visualizing collagen we used either second harmonic generated (SHG) images (Brown et al., 2007; Midgett et al., 2017; Sigal et al., 2014; Winkler et al., 2010) or polarized light microscopy (PLM) (Jan et al., 2017a, 2017b, 2015) or Instant polarized light microscopy (IPOLπ) (Lee et al., 2023). Neither of these techniques require labels or stains.

#### 2.3.1 Multicolor DiOlistics (MuDi) labeling

To visualize individual astrocytes within the LC we used multicolor DiOlistics (MuDi) labeling of vibratome sections as described in our previous publications (Waxman et al., 2025, 2023). We used gold microcarriers 1 µm in diameter coated with all combinations of 3 spectrally distinct dyes, DiI (Sigma-Aldrich 42364, 549/567 nm), DiO (Invitrogen D275, 484/501 nm), and DiD (Invitrogen D7757, 644/665 nm excitation/emission). Dye-coated microcarriers were loaded into Tefzel tubing (Bio-Rad, 1652441) and propelled into tissues using a Helios Gene Gun (Bio-Rad, 1652411) at 200 PSI. Microcarriers that contacted individual cells transferred dye to the cell membrane of those cells, allowing for visualization and distinguishing individual cells. Overall, this results in stochastic labeling of individual cells with one or more of the seven dye combinations. It is possible that not all cells labeled were astrocytes. However, we have shown that this technique ends up labeling a substantial number of individual cells that can be recognized as astrocytes based on their morphology (Waxman et al., 2025, 2023). Ultimately, our conclusions do not rest on the cells being astrocytes, but rather on them being cells that should not be within a blood vessel.

#### 2.3.2 DAPI staining

The vibratome sections and the cryosections were triple washed with PBS, each section was stained with 200 µl DAPI solution (Thermo Scientific, 0.1mg/µl, dilution 1:1000) for 10 minutes and triple washed with PBS again with PBS (Waxman et al., 2023).

#### 2.3.3 Immunostaining

For GFAP staining the vibratome sections were processed as previously described (Waxman et al., 2023). Briefly, sections were blocked and permeabilized in 5% BSA and 0.05% Triton X-100 for 24 hours, then incubated for 48 hours with a 1:200 GFAP primary antibody (rabbit, Invitrogen, PA5–16291). After a 24-hour PBS wash, they were incubated with a 1:500 secondary antibody for 24 hours, followed by three 10-minute PBS washes. Samples were processed through the same ethanol gradient, rehydrated for imaging, and mounted in a water-based medium with the imaged side facing up.

### 2.4 Imaging

#### 2.4.1 Imaging for collagen

SHG imaging is well-established for visualizing the collagen beam structure in the LC region. SHG images were acquired using an A1 Nikon multiphoton microscope (Tokyo, Japan). PLM is also a powerful technique for visualization and quantitative study of collagen fibers. As previously described, PLM images were acquired using a commercial inverted microscope (IX83, Olympus, Tokyo, Japan) with two polarized filters (Hoya, Tokyo, Japan), a polarizer placed before the sample and an analyzer after (Jan et al., 2017a, 2017b, 2015). IPOLπ is a recent version of PLM with several advantages for this work, compared with PLM and other versions of IPOL (Lee et al., 2023). First, the orientation-encoding color is cyclic every 180 degrees (π radians), whereas in other IPOL versions is 90 degrees (π/2). Most importantly, IPOLπ images have a bright background, whereas IPOL and most PLM often have dark backgrounds. Dark backgrounds are convenient to focus on the tissues of interest, but bright backgrounds allow seeing the context. In this case, IPOLπ shows both the birefringent collagen and shape and texture of neural tissues within the pores. The IPOLπ images were captured with an inverted microscope (Olympus IX83, Olympus, Tokyo, Japan) with a white light source, circular polarizer, polarization decoder, and color camera (DP74, Olympus, Tokyo, Japan), as previously described (Lee et al., 2023).

#### 2.4.2 Imaging for neural tissue

Images of MuDi label were acquired with an upright confocal microscope (FV1200, Olympus) using 20x magnification objectives within 24-48 hours of MuDi labeling. Images of GFAP staining were completed with the same confocal microscope at 20x magnifications, with the same X, Y, and Z image sizes as used for imaging of DiOlistics.

### 2.5 Definition and analysis of the pseudo-central retinal blood vessel

We evaluated visually each of the section images to determine if they had a CRBV-like structure. This was defined as a region within the lamina cribrosa with particularly thick collagen. In pig, sheep and goat this region was sometimes, but not always, connected to the inferior (or ventral) side of the canal (Figure 1). If they had one, and every eye did, we classified the CRBV-like structures into three types: fully pseudo-CRBV if there was evidence of cells other than erythrocytes within; partial pseudo-CRBV if some of the region appeared as vessel and other parts did not; CRBV if the structure only exhibited a blood vessel. Because mature erythrocytes do not have nuclei, we considered that observing nuclei within the CRBV-like structure indicated that this was not a vessel. The classification was conducted independently by a researcher who was blinded to the identity of the species. The data analysis and the chart were done in GraphPad (version 9.0).

## Results

We observed only true CRBVs at the center of every human and monkey eye, consistent with extensive literature (Albrecht May, 2008; Anderson, 1970, 1969; Hayreh, 1996) (Figure 1A, B). The collagen of the ONH in pig, sheep and goat exhibited similar CRBV-like structure (Figure 1C-E). We observed several pseudo-CRBVs with one side exhibiting neural tissue and another an opening with a blood vessel. We measured the CRBVs and pseudo-CRBVs areas in absolute magnitude and as a percentage of the LC canal area, and found no evidence that pseudo-CRBVs can be distinguished from CRBVs by their size (Supplementary Figure 1).

### 3.1 Cell Nuclei Presence Indicates Pseudo-CRBV

We observed cell nuclei presence within many CRBV-like structures, indicating pseudo-CRBVs. An example of goat is shown in Figure 2. With DAPI staining, we found cell nuclei presence inside the CRBV-like structure, indicating a pseudo-CRBV (Figure 2A-B). As per the definitions given in the methods, we considered this a partial pseudo-CRBV: the nuclear-rich regions were not vessels, and possibly were neural tissue. Importantly, there were also some blood vessels, that were absent of nuclei and had smooth thin walls (Figure 2C, blood vessel outlined in cyan). Inside the pseudo-CRBV, a strong collagen beam crosses the region, as shown in the SHG image in Figure 2A. In the DAPI staining of the same sample, cell nuclei along this collagen beam appear elongated and spindle-shaped, resembling fibroblasts (Figure 2D, yellow arrow). In contrast, cell nuclei in other areas within the pseudo-CRBV are more rounded, like those found in other pores of the ONH, suggesting they might be neural tissue (Figure 2D, blue arrow).

**Figure 2:**
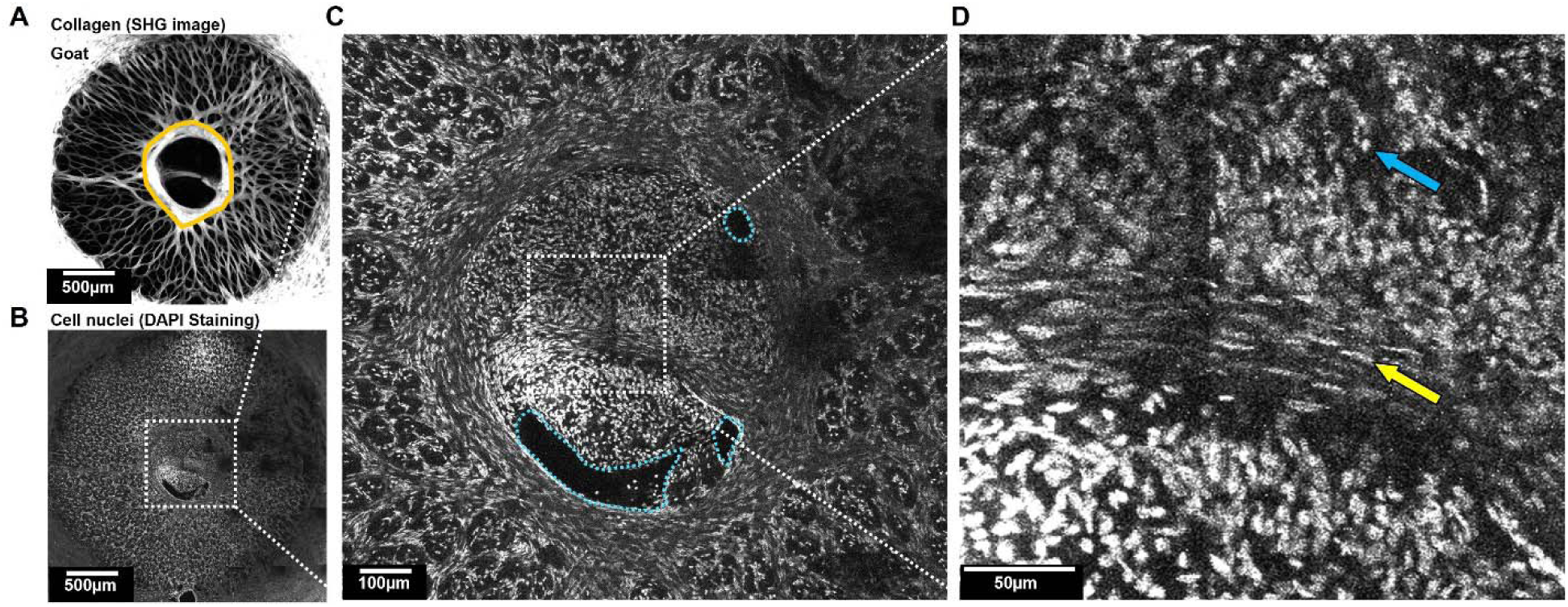
Cell Nuclei Presence Indicates Pseudo-CRBV. A) SHG image of goat LC showing the collagen network, with the CRBV-like structure outlined in orange. Notably, a strong collagen beam structure extends across the entire CRBV-like structure. This could have been interpreted as separating the central retinal artery and vein. B) This is the DAPI staining of the same goat section. The CRBV-like structure contains cell nuclei. C) Further zoom-in, we considered this a partial pseudo-CRBV: the nuclear-rich regions likely are neural tissue, while the nuclei-absent regions with smooth, regular walls were blood vessels, as outlined in cyan. D) The nuclei along the strong collagen beam appear elongated and spindle-shaped, resembling fibroblasts (yellow arrow). In contrast, cell nuclei in other areas within the pseudo-CRBV were more rounded, like those found in other pores of the ONH, suggesting they might be neural tissue (blue arrow).

### 3.2 Astrocyte Presence Indicates Pseudo-CRBV

#### 3.2.1 Multicolor DiOlistics Labeling Evidence

We found cells with astrocyte-like morphology inside the CRBV-like structure, indicating pseudo-CRBVs. MuDi labeling showed that true CRBVs in monkey lack cells within (Figure 3A). In contrast, in goat and sheep, MuDi-labeled cells were present inside the CRBV-like structure, indicating a pseudo-CRBV (Figure 3B-C). The pseudo-CRBV was composed of a varied population of cells, including astrocyte-like cells. These astrocyte-like cells had a single soma with multiple branches that made contact with the collagen structure, as described before (Waxman et al., 2025, 2023).

**Figure 3:**
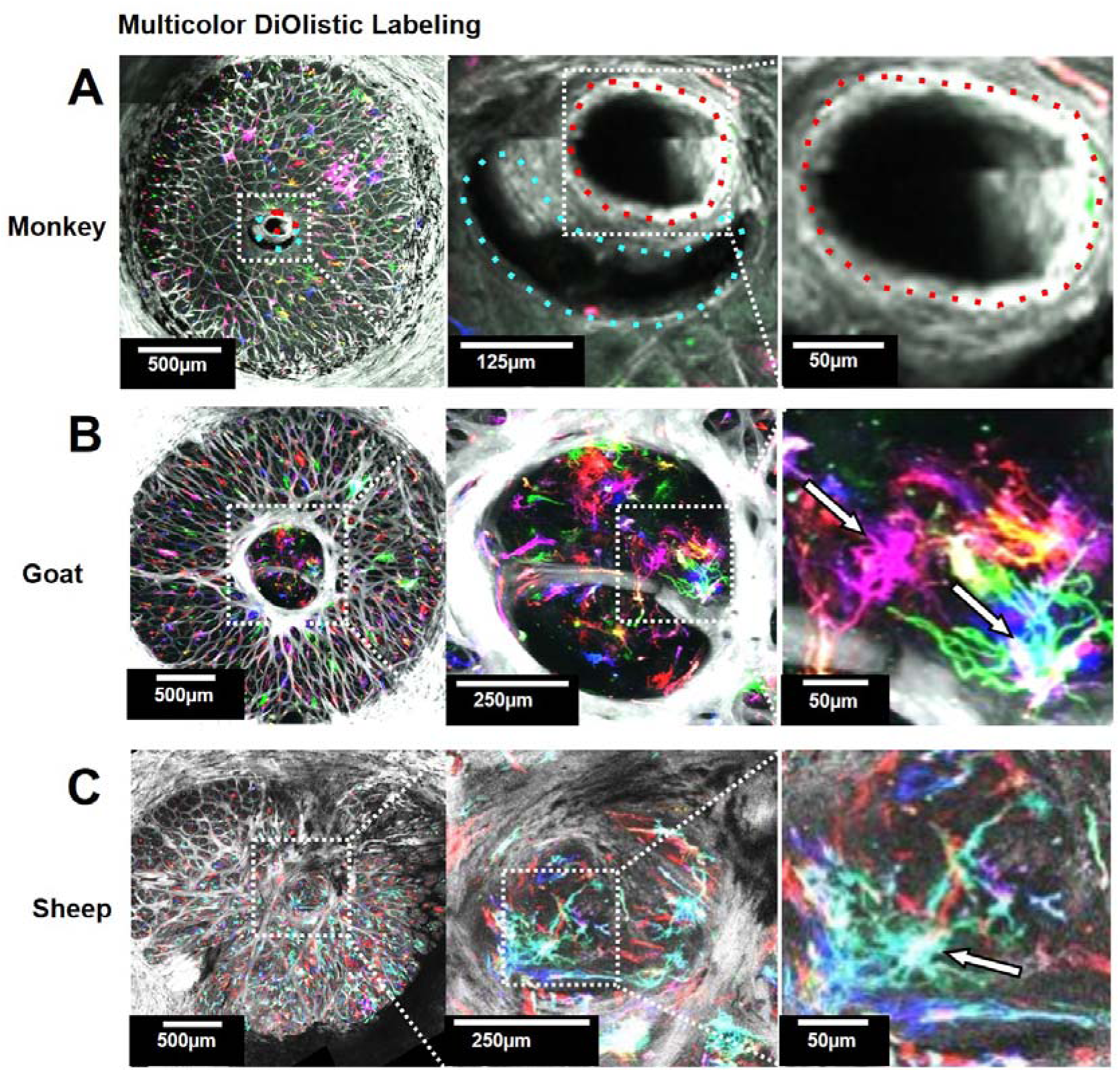
Cells with Astrocyte-like Morphology Presence Indicates Pseudo-CRBV. A) MuDi labeled confocal image of the monkey LC, showing that the real vessels do not contain any MuDi labeled cells. B) Merged SHG and MuDi labeled image of the goat LC. Inside the CRBV-like structure, it consists of a various population of cells and especially cells with astrocyte-like morphologies (white arrow). C) MuDi labeled confocal image of the sheep LC, showing the CRBV-like structure is filled with MuDi labeled cells. It consists of a various population of cells and cells with astrocyte-like morphologies (white arrow) as well.

#### 3.2.2 GFAP Immunostaining Evidence

We observed GFAP-positive astrocytes within the CRBV-like structure, again indicating pseudo-CRBVs. We utilized GFAP immunostaining and autofluorescence to visualize the astrocytes and collagen, respectively. Where CRBVs would be expected, a collagen structure is visible, outlined in orange (Figure 4A). This region, however, predominantly contained astrocytes. There were a couple of small openings, outlined in cyan, that could have been vessels (Figure 4B-C). This finding suggested that astrocytes were a key component of the pseudo-CRBV, contributing to their cellular makeup.

**Figure 4:**
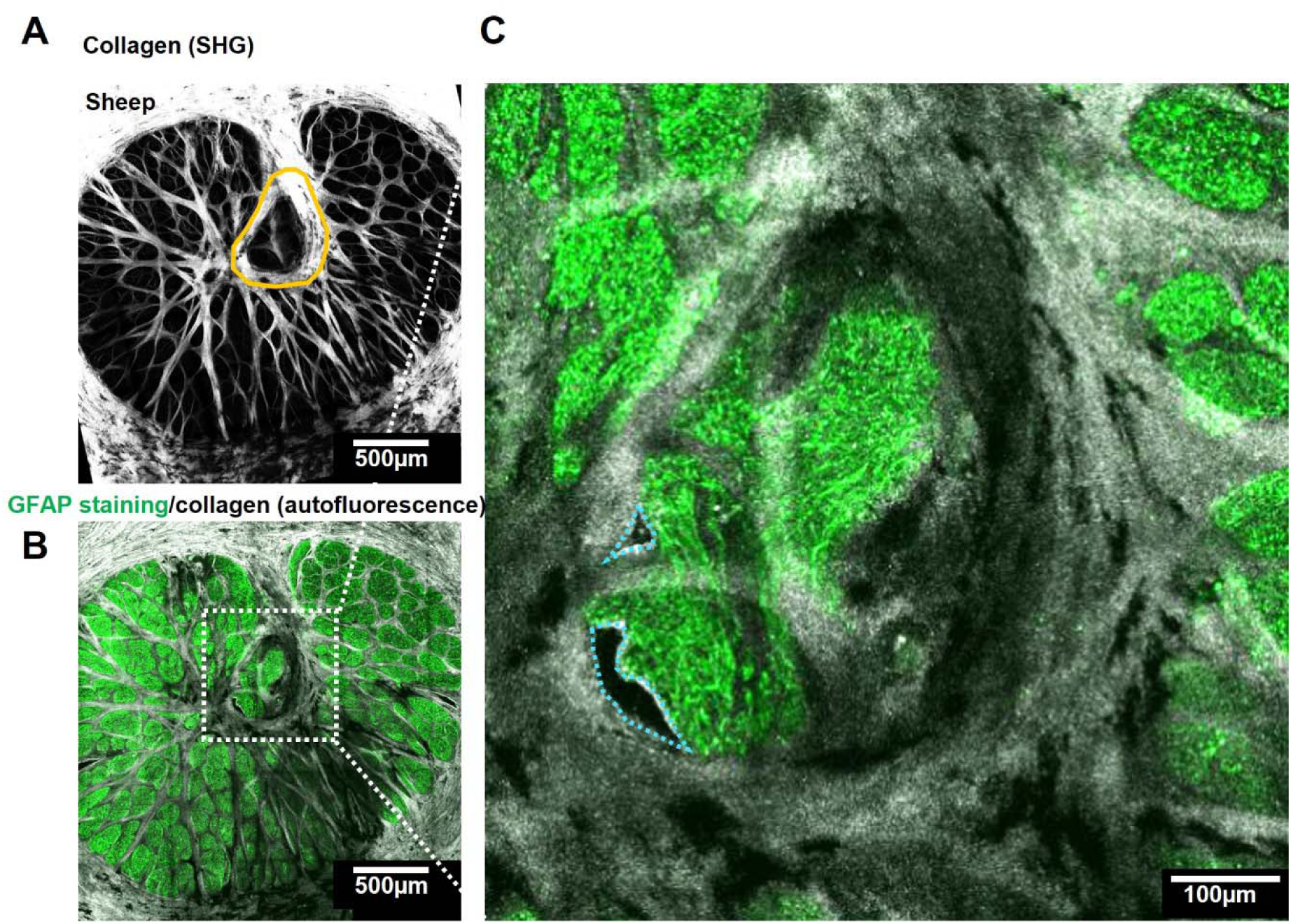
Astrocytes Presence Indicates Pseudo-CRBV. A) SHG image of a sheep LC. A CRBV-like structure is outlined with an orange line. Notice the relatively thick collagen walls near the central LC. B) The same section after GFAP staining and autofluorescence imaging with a confocal microscope. The CRBV-like structure is shown to contain astrocytes over most of the lamina. C) Detail reveals mainly coverage of this region with astrocyte GFAP. A small portion of this region could be occupied by blood vessels, outlined in cyan. Hence, this example was classified as a partial CRBV.

### 3.3 More than Half of the CRBV-like structures in Pig, Sheep and Goat were Pseudo-CRBV

To strengthen our argument, we evaluated labeling vibratome slabs (150 um-thick) using Lycopersicon esculentum (Tomato) lectin (Invitrogen, L32472), as described in previous paper (Bowden et al., 2025). An example pig LC labeled this way and imaged using a multiphoton microscope is shown in Figure 5. The CRBV-like structure shows a couple of small capillaries but is still clearly a pseudo-CRBV.

**Figure 5:**
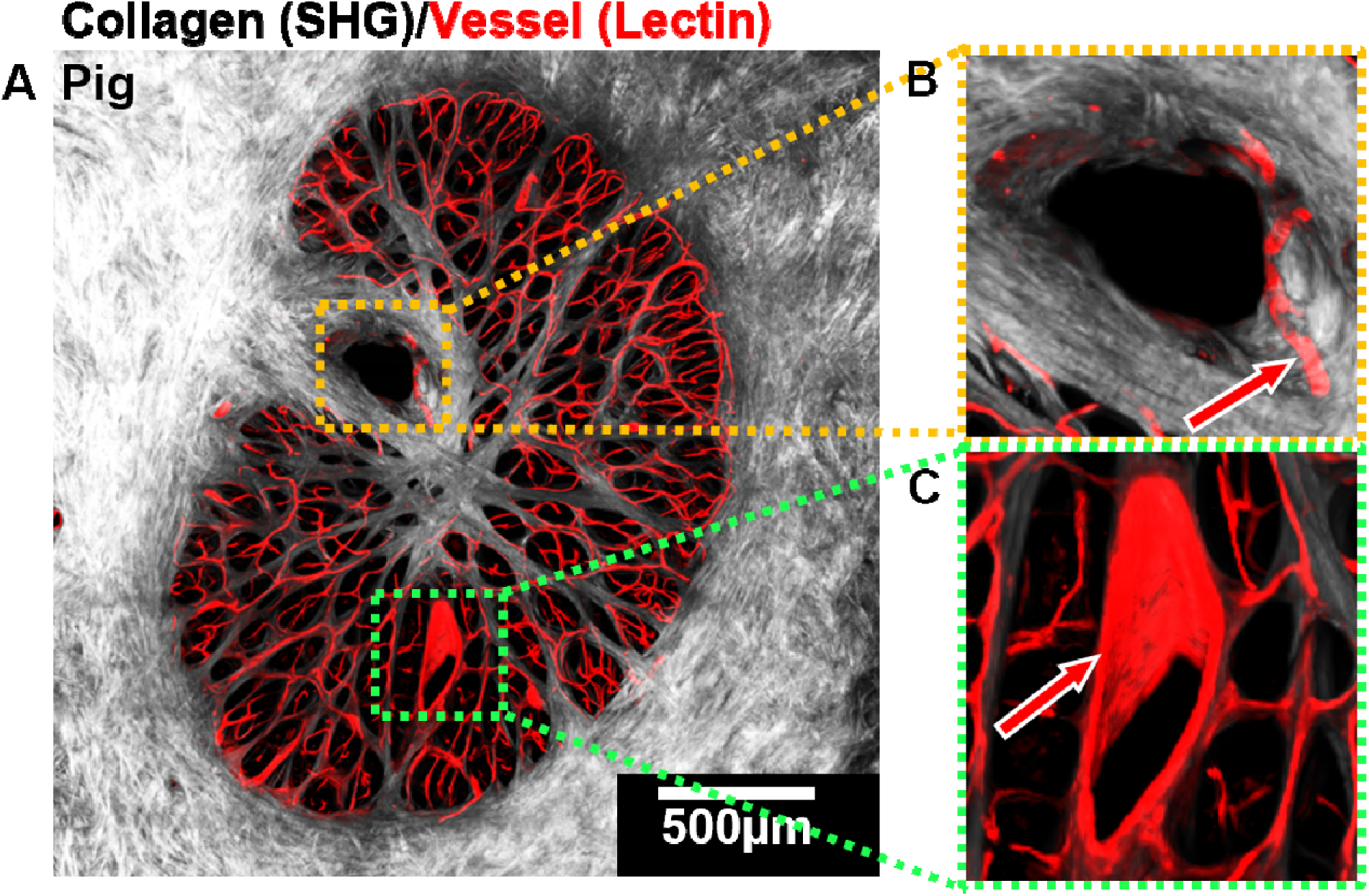
Example of pseudo-CRBV in pig eye with vessels labeled. Collagen, visualized by second-harmonic generation (SHG), is shown in gray, and tomato lectin-labeled blood vessels are shown in red. A) Overview of the entire optic nerve head canal. B) Close-up view of the CRBV-like structure, showing the absence of major lectin-labeled vessels inside. C) Close-up view of a large vessel with a lectin-positive wall located outside the CRBV-like structure. The presence of lectin-positive small vessels in (B) and the large vessel in (C) demonstrates successful lectin labeling in this sample. Together, these observations support that this pseudo-CRBV region is negative for lectin, consistent with other observations in this paper.

Figure 6A-C illustrates the process to distinguish CRBV-like structures. The process included assembling images from several consecutive 16-µm cryosections with DAPI staining merged with IPOLπ in pig and sheep. This was crucial to rule out that potential artifacts in one section could give the wrong result. In goat eyes, we used vibratome sections and used different labels to visualize the content inside CRBV-like structure to determine a pseudo-CRBV or not. We found that in pig eyes, 20% (1/5) of eyes were CRBVs, 60% (3/5) were partial pseudo-CRBVs and 20% (1/5) were fully pseudo-CRBVs (Figure 5D). In sheep eyes, 50% (3/6) of eyes were CRBVs, 33.3% (2/6) were partial pseudo-CRBVs, and 16.7% (1/6) were fully pseudo-CRBVs (Figure 5D). In goat eyes, 100% (2/2) were partial pseudo-CRBVs. Overall, more than half of the CRBV-like structures were pseudo-CRBV in pig (80%), sheep (50%) and goat (100%).

**Figure 6:**
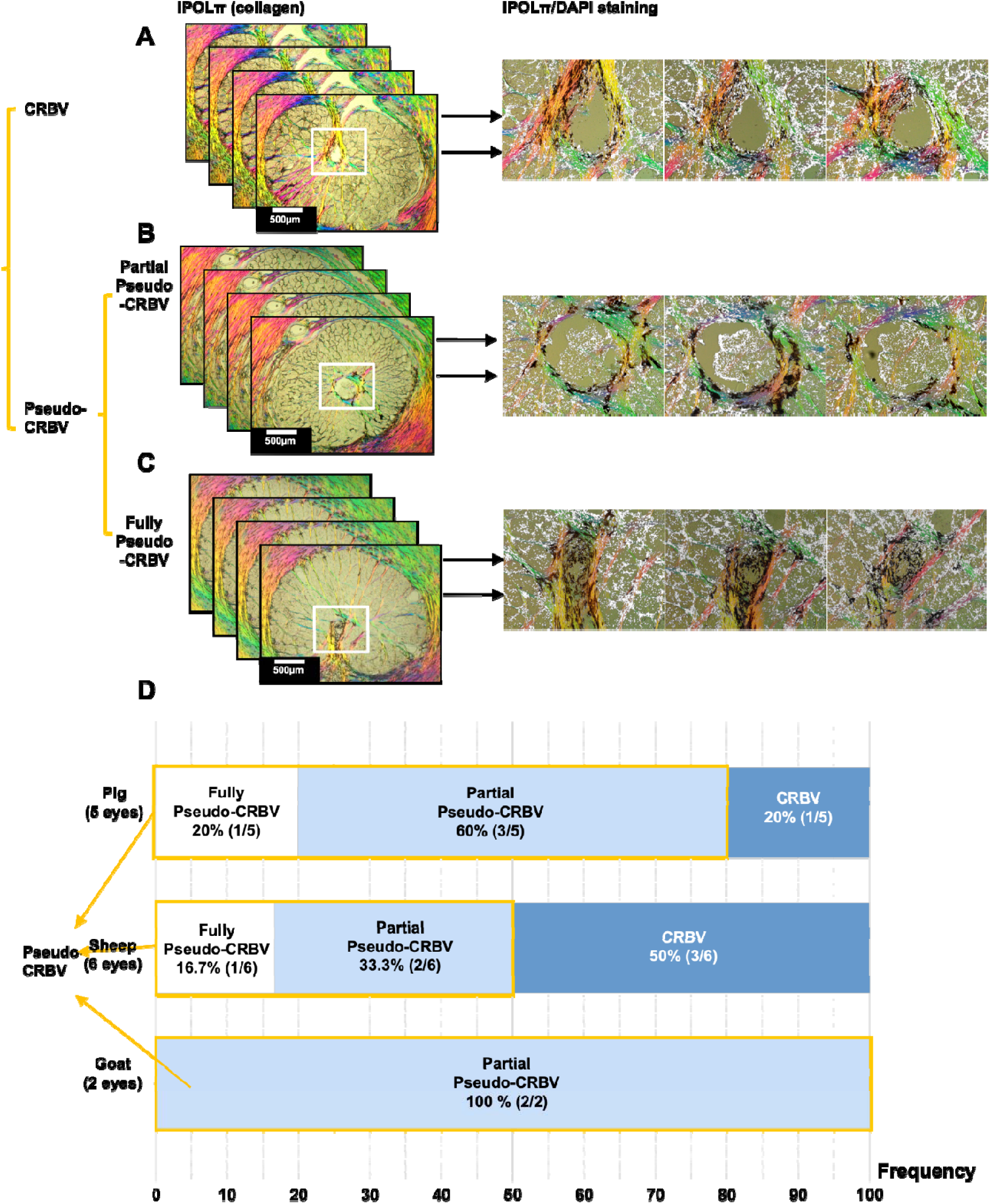
Example of CRBV-like classification. (A-C) Three stacks of 16 µm-thick cryostat sections captured by IPOLπ. To avoid potential artifacts in single sections, the classification was based on multiple sections. In this case three sections are shown for each stack. The images show merged DAPI and IPOLπ images, which allows discerning the nuclei (white) relative to the collagen (colors) and pigment (black). Colors in IPOLπ indicate local collagen orientation. The three stacks were selected as examples of each of the types of CRBV-like structures: 1. A CRBV if the evidence was consistent with the structure functioning as a blood vessel. 2. Partial pseudo-CRBV if the evidence applied only on part of the region, while other regions still appear as vessels; 3. Fully pseudo-CRBVs if there was evidence of neural tissues or other cells within (other than RBCs); D-E) Bar chart of the frequency of each type. More than half of the CRBV-like structures are pseudo-CRBV in pig (80%), sheep (50%), and goat (100%).

## Discussion

Our goal was to determine if the CRBV-like structures in pig, sheep and goat ONHs are true CRBVs, or if they should, instead, be considered pseudo-CRBVs. The results demonstrate that in these three species, the structures frequently contain neural tissue and non-vascular cells and therefore cannot always be true vessels. Pseudo-CRBVs occurred in the majority of the eyes examined – 80% (4/5) in pig, 50% (3/6) in sheep and 100% (2/2) in goat, indicating that relying on the collagen to identify CRBVs may lead to misinterpretations. These results contrast with human and monkey eyes in which the structures are always CRBVs. Below we discuss the findings and their implications in more detail.

### 4.1 Evidence Supporting the Identification of the Pseudo-CRBV

Several lines of evidence support our proposal to consider these structures as pseudo-CRBVs. First, DAPI staining often revealed nuclei within the region. Because mature erythrocytes lack nuclei, this finding is inconsistent with a vessel lumen. Second, MuDi labeling and GFAP immunostaining demonstrated the presence of astrocytes and other cells with the morphology of neural cells, within the regions. Altogether, vascular structures were occasionally observed, the regions appearing vessel-like in the images of collagen more often were not large vessels.

We considered whether our observations could result from artifacts, such as sectioning distortions or projection effects. We examined carefully the sections for folds or tissue breaks, but did not see anything that could explain the observations. The consistency of the findings across multiple serial sections of several eyes, in thin and thick sections, and through imaging modalities argues strongly against this possibility. Although in some eyes CRBV-like regions did contain vessels, most often these were relatively small ones, not the large-caliber consistent with a central retinal artery and vein pair observed in the human and monkey (Albrecht May, 2008; Anderson, 1970, 1969; Hayreh, 1996).

SHG, PLM and IPOL imaging are well-established techniques for visualizing the collagen structure in the ONH and other tissues (Behkam et al., 2019; Brown et al., 2007; Czerpak et al., 2023; Ling et al., 2019; Midgett et al., 2020, 2017; Pijanka et al., 2019; Sigal et al., 2014; Winkler et al., 2010). Although the images showed CRBV-like structures in every eye studied, this was based on a qualitative and subjective interpretation. Given the fundamental functional and microstructural differences between the collagen of vessels and the lamina cribrosa beams, it seems plausible that closer analysis of the collagen images or labeling for specific collagen types, may reveal whether a CRBV-like structure on first look is an actual vessel or a pseudo-CRBV.

MuDi labeling can reveal the cell morphologies of neurons and astrocytes in central nervous system tissues (Foster Olive et al., 2018; Hough and Brown, 2017; O’Brien and Lummis, 2006; Seabold et al., 2010; Staffend and Meisel, 2011). In our previous work, we used MuDi labeling to show the individual astrocytes and their morphologies in the LC region (Waxman et al., 2025, 2023). In this work, we identified cells with astrocyte-like morphology within the CRBV-like structure, indicating a pseudo-CRBV. Some cells exhibited a single soma with branching processes, characteristic of astrocytes. Further GFAP staining confirmed the presence of astrocytes within the pseudo-CRBV.

### 4.2 If not vessels, what are the pseudo-CRBVs?

Our findings raise questions regarding the origin and function of pseudo-CRBVs. If not vessels, what are these structures? What is their function and why does their collagen appear similar to that of actual CRBVs? What process underlies their formation such that some of them contain actual vessels whereas others do not? While we do not know the answers, we would like to propose two potentials, not mutually exclusive, explanations: one biomechanical, the other developmental.

<u>Biomechanical explanation:</u> in humans and monkeys the CRBVs provide important, potentially critical, structural support to the ONH and its ability to bear the loads of intraocular and cerebrospinal fluid pressures and of gaze-induced deformations (Choung et al., 2021; Feola et al., 2018; Hua et al., 2018; Huang et al., 2013; Otani et al., 2024; Panda et al., 2022; Sawada et al., 2021a, 2021b; Sigal et al., 2005; Wang et al., 2017, 2016; Wei et al., 2022a). The position of the central retinal vessel trunk within the ONH has been associated with the pattern of visual field loss and the thickness of the retinal nerve fiber layer in advanced stages of glaucoma (Budde and Jonas, 2003; Huang et al., 2013). Specifically, a more centrally located vessel trunk was associated with less neural tissue damage. Additional evidence indicates that blood pressure, and the associated hydraulic-stiffening of the CRBVs contribute to provide structural support to the optic nerve head (Tran et al., 2018; Wei et al., 2022b). We posit that pig, sheep and goat may have similar biomechanical requirements, and therefore these species have developed CRBV-like collagen structures for biomechanical reasons, namely for structural stability and to help bear loads even if these are not directly tied to perfusion needs.

<u>Developmental explanation:</u> Another possible explanation is that the pseudo-CRBVs remnants of the embryonic hyaloid vasculature. In humans, the hyaloid artery (HA) is a transient network of fetal vasculature formed during ocular development that delivers oxygen and nutrients to the embryonic eye. The HA enters the eye from the choroid fissure in the optic cup, passes through the vitreous and contacts the posterior pole of the lens. It is important for embryonic development of the lens, vitreous and retina. The HA undergoes developmental regression as soon as the retinal vasculature starts to form (Saint-Geniez and D’Amore, 2004). The developmental fate of the hyaloid vasculature in other species, however, remains less well characterized. To our knowledge, only one study has suggested that the hyaloid vasculature in pigs is distinct at 4 weeks of gestation, begins to regress by 8 weeks, and undergoes complete regression during the early postnatal period (De Schaepdrijver et al., 1989).

The CRBV-like structures in pig, sheep, and goat may be related to the hyaloid vasculature in forming and extending into the vitreous body. These collagenous structures may persist beyond the regression of the hyaloid in later phases of development. Interestingly, in several eyes, the pseudo-CRBVs structures appear to contained neural tissues, the reason for which remains unclear. One possible explanation is that neural tissue grows into these collagenous remnants during a similar developmental time window as hyaloid vessel regression.

To us, the non-collagen components of the pseudo-CRBVs, i.e. what appears as the lumen of a CRBV, often resembled other LC pores: they have neural tissues, including axons and astrocytes, and small vessels for local perfusion. These small vessels were capillaries, too small and local to be responsible for central retinal supply or drainage. Nevertheless, whether the pseudo-CRBVs are distinct from other pores in terms of structure or function remains unknown.

### 4.3 Pseudo-CRBV is Beyond Individuals in Pigs, Sheep, and Goat

Another interesting aspect of our results is that in some eyes the CRBV-like structures were indeed CRBVs. The mechanisms underlying why some species only have actual CRBVs and others have both CRBVs and pseudo-CRBVs remain unknown. The variations could be a consequence of the structure roles, as discussed above, or they could be due to differences in the overall vascular supply among species. Pigs, sheep and goat are thought to rely more on ciliary circulation for posterior pole perfusion than humans and monkeys which rely heavily on the central retinal vein (Rootman, 1971). In pigs, up to six branches of retinal arteries enter the ONH at the level of the sclera, with the retinal veins forming only one CRV that exits the eye through the LC region(Simoens et al., 1992). Limited anatomical information is available for the analogous region in sheep.

### 4.4 Limitations

We must acknowledge several limitations in this work. First, the distribution of the pseudo-CRBV was based on the eyes of 5 pigs, 6 sheep, and 2 goats. While the sample size is small and the percentages may not fully reflect the true distribution across all animals from these species, the sample size is sufficient to recognize that a significant proportion of the eyes from these species are pseudo-CRBVs. We have to acknowledge the possibility that what we report herein is not general for the species we studied, and that particular breeds, subspecies or strains vary. We were able to confirm that the pig eyes obtained from Thoma Meat Market in Saxonburg PA were from Yorkshire white pigs. The specific breeds of the sheep and goat eyes procured through MRG Food, McKeesport, PA were not available. Second, the staining and imaging techniques we used do not provide a complete view of the cells and tissue components and their architecture. For example, DAPI revealed cell nuclei, but does not reveal cell shape or type. In addition, within the complex ONH architecture it is possible that in some sections we could have misidentified an actual CRBV as a partial pseudo-CRBV. This possibility seems less likely given that we also observed pseudo-CRBVs in 3D visualizations. A more comprehensive study fully mapping in 3D the various components of the ONH, including vessels, collagen, astrocytes, axons, etc. would help improve the tissue characterization. Further research looking into the eyes of other species, such as cow or dog, could help to better understand the extent to which pseudo-CRBVs are a widespread phenomenon on large mammals or is it potentially limited to the three species studied in this work.

In conclusion, to the best of our knowledge, this study provides the novel insight that CRBV-like structures in pig, sheep and goat eyes are often, but not always, pseudo-CRBVs. The mechanisms leading to this situation and the consequences remain unknown.

## Notes

This work was supported in part by the National Institutes of Health R01-EY023966, P30-EY008098, and T32-EY017271 (Bethesda, MD). The Eye and Ear Foundation (Pittsburgh, PA). Research to Prevent Blindness (unrestricted grant to UPMC ophthalmology, and Stein innovation award to I.A. Sigal) and the Wiegand Fellowship in Ophthalmology (to S. Waxman).

### Competing Interest Statement

The authors have declared no competing interest.

