## Supplementary Figure 1 for "Pseudo-Central Retinal Blood Vessel in Pig, Sheep and Goat"

**Supplementary Material**

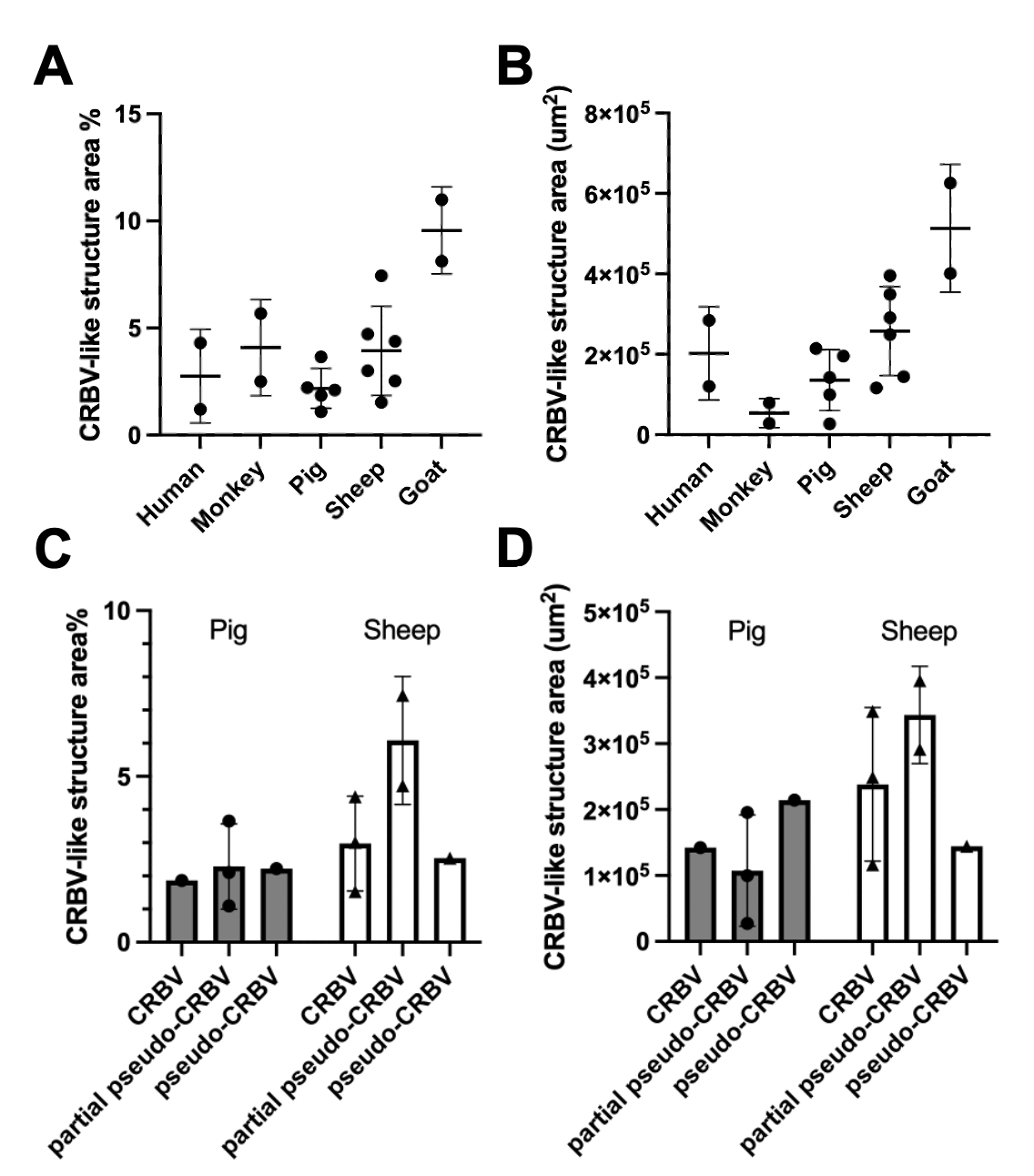
 **Supplementary Figure 1 Quantification of the CRBV-like structure area across species and within species.** We measured the CRBVs and pseudo-CRBVs areas in absolute magnitude and as a percentage of the LC canal area. Because of the small number of samples, we did not do statistical analyses. Nevertheless, from inspection it seems like there were differences between species in the size of the CRBV-like structures. Monkeys had the smallest and goats the largest. The plots on the bottom row show the sizes of the pig and sheep CRBV-like structures split by category. Overall these results do not support the conclusion that the pseudo-CRBVs are any larger, or smaller, than CRBVs.
